# Nardilysin fine-tunes the mammalian circadian clock through selective modulation of PER2 function

**DOI:** 10.64898/2026.08.17.745167

**Authors:** Yoshinori Hiraoka, Rina Nunokawa, Mikiko Ohno, Yusuke Morita, Yuki Kato, Kiyoto Nishi, Noriaki Kume, Yoshitaka Fukada, Hikari Yoshitane, Eiichiro Nishi

## Abstract

Circadian rhythms in mammals are generated by negative feedback loops, in which CLOCK and BMAL1 bind to E-box to activate transcription of *Period* (*Per*) and *Cryptochrome* (*Cry*) and the E-box-dependent transactivation is inhibited by PER and CRY proteins. Although the core transcriptional feedback loop of the circadian clock has been well defined, how this machinery interfaces with broader nuclear regulatory systems remains incompletely understood. Here, we identify nardilysin (NRDC), a metalloendopeptidase previously implicated in nuclear transcriptional regulation and metabolic homeostasis, as an unexpected modulator of the circadian clock. NRDC deficiency led to elevated PER2 protein levels in the liver and enhanced PER2 dynamics in cell-autonomous circadian oscillators, and was accompanied by a significant shortening of behavioral rhythms in mice. Biochemical analyses demonstrated that NRDC selectively associates with PER2 and CRY2 and antagonizes PER2-mediated repression of CLOCK–BMAL1–dependent transcription. Genome-wide chromatin immunoprecipitation analyses reveal that NRDC is enriched at promoter-proximal E-box-containing regions, frequently co-localizing with CLOCK binding sites. Together, these findings uncover a previously unrecognized link between circadian timing and protease-based nuclear regulation, positioning NRDC as a critical modulator of PER2 function and circadian period determination.

## Introduction

The circadian clock is a timing system that enables organisms to align biochemical, physiological, and behavioral rhythms in resonance with the 24-hour environmental cycles. In mammals, the molecular clock is composed of transcriptional/translational negative feedback loops (TTFLs) (1–3). The transcription factors CLOCK and BMAL1 activate transcription of *Period* (*Per*) and *Cryptochrome* (*Cry*) via E-box elements. The translated PER (PER1, PER2, PER3) and CRY (CRY1, CRY2) proteins accumulate, assemble into a large nuclear PER complex (4, 5), and repress their own expression by binding to the CLOCK-BMAL1 complex (6–8). Turnover of PER and CRY proteins (9–12) then permits reactivation of the E-box-dependent transcription, initiating the next cycle. This core oscillator drives rhythmic expression of a wide array of downstream genes, thereby coordinating diverse physiological processes including metabolism, immune responses, and body temperature regulation (13–15).

While the canonical components of the circadian clock have been extensively characterized, the system is further modulated by multiple regulatory layers that integrate chromatin dynamics, post-translational modification, and proteolytic control. These factors fine-tune circadian timing and amplitude, ensuring both robustness and adaptability of the oscillator. For example, phosphorylation of PER proteins by casein kinase 1 (CK1), a PER-interacting kinase, regulates PER protein stability and nuclear localization, thereby determining clock oscillation speed (16–18). Other PER-interacting proteins such as CAVIN-3 and promyelocytic leukemia (PML), also modulate PER2 function and clock regulation (19, 20). Several RNA-binding proteins and chromatin modifying factors have also been identified as constituents of nuclear PER complexes (4, 5, 21, 22), where they recruit SIN3 histone deacetylase (HDAC) (21) and SUV39H histone methyltransferase (HMT) complexes (22) to augment the PER-mediated transcriptional repression.

Nardilysin (N-arginine dibasic convertase; NRDC) is a zinc peptidase of the M16 family, which selectively cleaves dibasic sites (23, 24). While NRDC was originally characterized for extracellular functions that enhance ectodomain shedding of membrane proteins (25–30), subsequent studies have revealed a prominent nuclear role for NRDC in transcriptional regulation. NRDC preferentially binds H3K4me2 on histone H3 tails and associates with the NCoR/SMRT/HDAC3 corepressor complex at target genes (31). It also regulates diverse homeostatic programs through interactions with transcriptional modulators, including PGC-1α in brown adipocytes, Islet-1 in pancreatic β-cells, and HDAC1/p53 in intestinal epithelial cells (32–34). Consistent with these nuclear activities, chromatin immunoprecipitation-sequencing (ChIP-seq) in immortalized MEFs showed that NRDC helps maintain an appropriate epigenetic landscape and supports orderly cell-cycle progression (35).

Accumulating evidence indicates that the circadian clock is deeply integrated with diverse physiological programs that maintain cellular and systemic homeostasis (2, 13–15). Beyond its role in generating daily rhythms, the clock interfaces with nuclear transcriptional networks that coordinate metabolism, inflammatory responses, and tissue-specific gene expression. NRDC has been repeatedly implicated in such homeostatic regulation, acting as a metalloendopeptidase with prominent nuclear functions in transcriptional control across multiple biological contexts, including metabolism and inflammation (32–34). Given its ability to associate with chromatin-related complexes and modulate transcription through protein–protein interactions, NRDC represents a compelling but previously unexplored candidate for linking circadian timing to broader nuclear regulatory systems. Whether and how this protease-based nuclear regulation intersects with the core circadian feedback loop has remained unknown.

## Results

### NRDC deficiency shortens the circadian period *in vivo*

To examine whether NRDC’s diverse homeostatic regulation extends to modulate the mammalian circadian rhythm, we first monitored free running activities of wild-type (WT) and NRDC whole body knockout (NRDC-KO) mice (29) by using an area sensor. While both WT and NRDC-KO mice exhibited robust activity rhythms in LD and DD (Fig. 1A), the circadian period of NRDC-KO mice was significantly shorter than that of WT mice under DD conditions (23.83 ± 0.16 h in WT versus 23.56 ± 0.14 h in NRDC-KO) (Fig. 1B). Importantly, we previously demonstrated that NRDC is a key molecule in determining the set point of body temperature and that NRDC-KO mice exhibited an approximately 1.5 °C lower body temperature under our housing condition (23 ± 1 °C) (32). A fundamental property of the circadian clock is called as temperature compensation, whereby the period length remains largely unchanged despite fluctuations in ambient temperature, whether in summer or winter. In mammals, however, brain temperature is maintained within an extremely narrow range under physiological conditions, making it difficult to directly assess the functional significance of temperature compensation *in vivo*. In this study, we took advantage of NRDC-deficient mice, in which brain temperature is measurably reduced. Strikingly, despite the lowered brain temperature, the circadian clock did not slow down as might be expected from basic biochemical kinetics. Instead, we observed a tendency toward period shortening. These findings provide *in vivo* evidence for the remarkable robustness of temperature compensation in the mammalian circadian system. Furthermore, they suggest that NRDC actively contributes to the regulation of the circadian period *in vivo*, rather than merely influencing body temperature as a secondary effect.

**Fig. 1.**
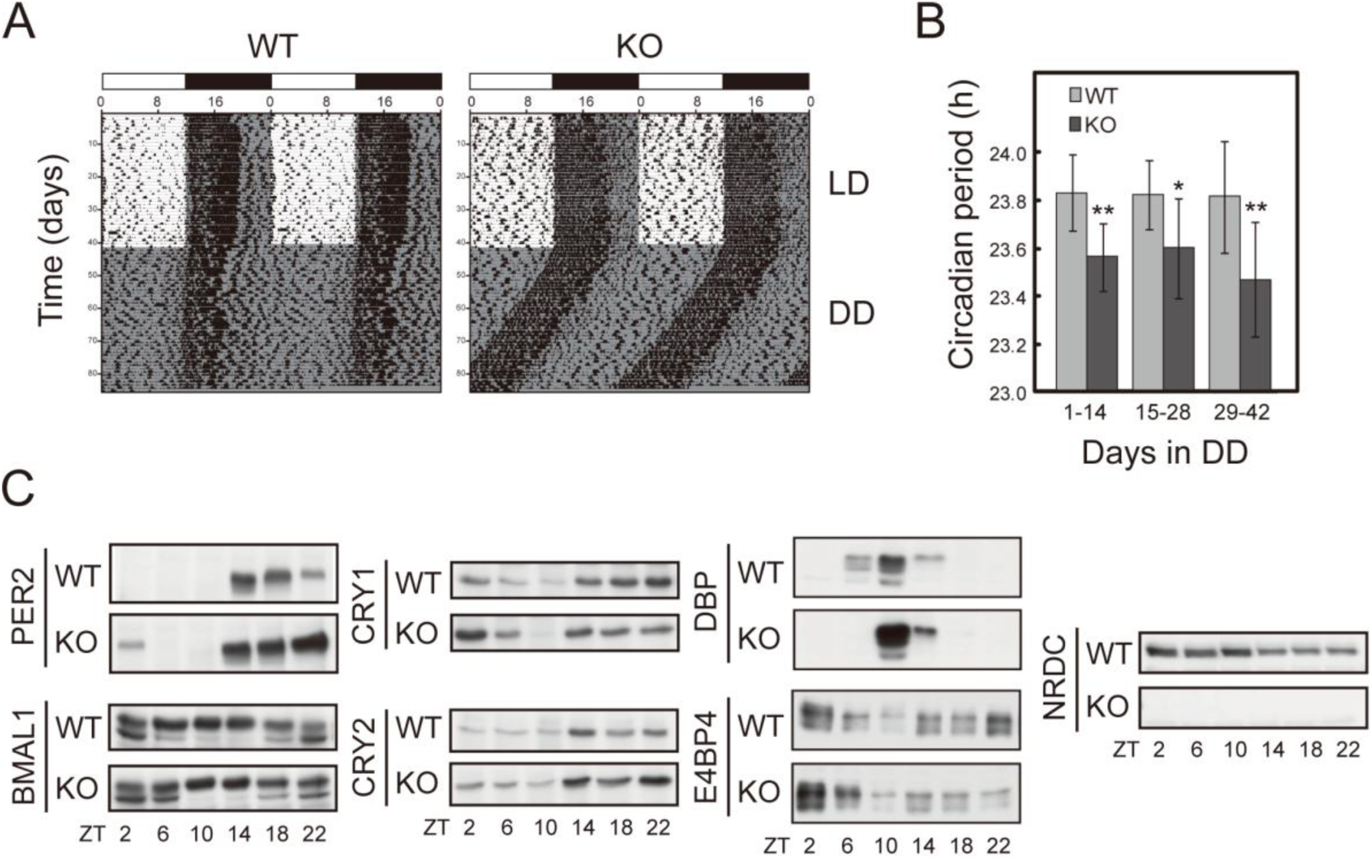
Shortened circadian behavioral rhythms and altered circadian expression of clock proteins in NRDC-deficient mice. (A) Representative actograms of the activities of WT and NRDC-KO mice. Mice were entrained to 12-h light/12-h dark (LD) cycle for 6 weeks and then transferred to constant darkness (DD). White and black bars indicate the light and dark phases in LD cycle, respectively. (B) Circadian periods of the behavior rhythms under DD condition in WT and NRDC-KO mice. Data are presented as mean ± SD (*n* = 10 for WT; *n* = 9 for NRDC-KO. \**P* < 0.05; \*\**P* < 0.01). (C) Circadian accumulation of clock proteins in nuclear extracts from WT and NRDC-KO mouse livers collected at 4-h intervals. Immunoblotting was performed with the indicated antibodies.

### NRDC affects circadian expression of core clock proteins and PER2::Luc responses

To assess the role of NRDC in the circadian molecular oscillation, we examined the circadian expression profiles of NRDC and core clock components in the liver at protein levels (Fig. 1C). Due to their reduced body temperature, NRDC-KO mice are prone to postnatal cannibalization, which substantially limits the number of animals available for study (approximately 80% died within 48 h of birth) (29). Nevertheless, we collected a sufficient number of KO mice to assess the temporal profile at six time points. Then, WT and NRDC-KO mice were maintained under DD conditions, and liver samples were collected at multiple Zeitgeber times across the circadian cycle. NRDC protein levels remained relatively constant throughout the day, with a modest reduction during the dark phase (Fig. 1C). The NRDC signals were completely abolished in NRDC-KO livers. Temporal profiles of clock proteins have apparent changes, especially PER2, CRY1 and E4BP4 at ZT2 and DBP at ZT10, while the number of biological replicates was constrained by the limited availability of KO mice (Fig. 1C).

We next examined whether NRDC modulates the cell-autonomous circadian oscillator using mouse embryonic fibroblasts (MEFs) derived from WT and NRDC-KO mice carrying the PER2::LUC knock-in allele. In these cells, a PER2-luciferase fusion protein is expressed under the control of endogenous *Per2* promoter (36). PER2 induction in response to serum shock (37) was significantly higher in the NRDC-KO MEFs (Fig. S1), suggesting that NRDC may help restrain excessive feeding-induced increases in PER2 protein levels. This interpretation is consistent with the increased nocturnal PER2 protein levels in NRDC-KO liver (Fig. 1C).

### NRDC interacts with PER2 and CRY2 through defined protein domains

We next examined the interaction of NRDC with core components of the circadian feedback loop, including CLOCK, BMAL1, PER2, and CRY2, in HEK293T/17 cells. We co-expressed NRDC-V5 with myc-CLOCK, FLAG-BMAL1, HA-PER2, or HA-CRY2, and performed co-immunoprecipitation using anti-V5 antibody. This revealed that both PER2 and CRY2 were co-precipitated with NRDC-V5 (Fig. 2 A and B), whereas CLOCK and BMAL1 were not (Fig. S2 A and B).

**Fig. 2.**
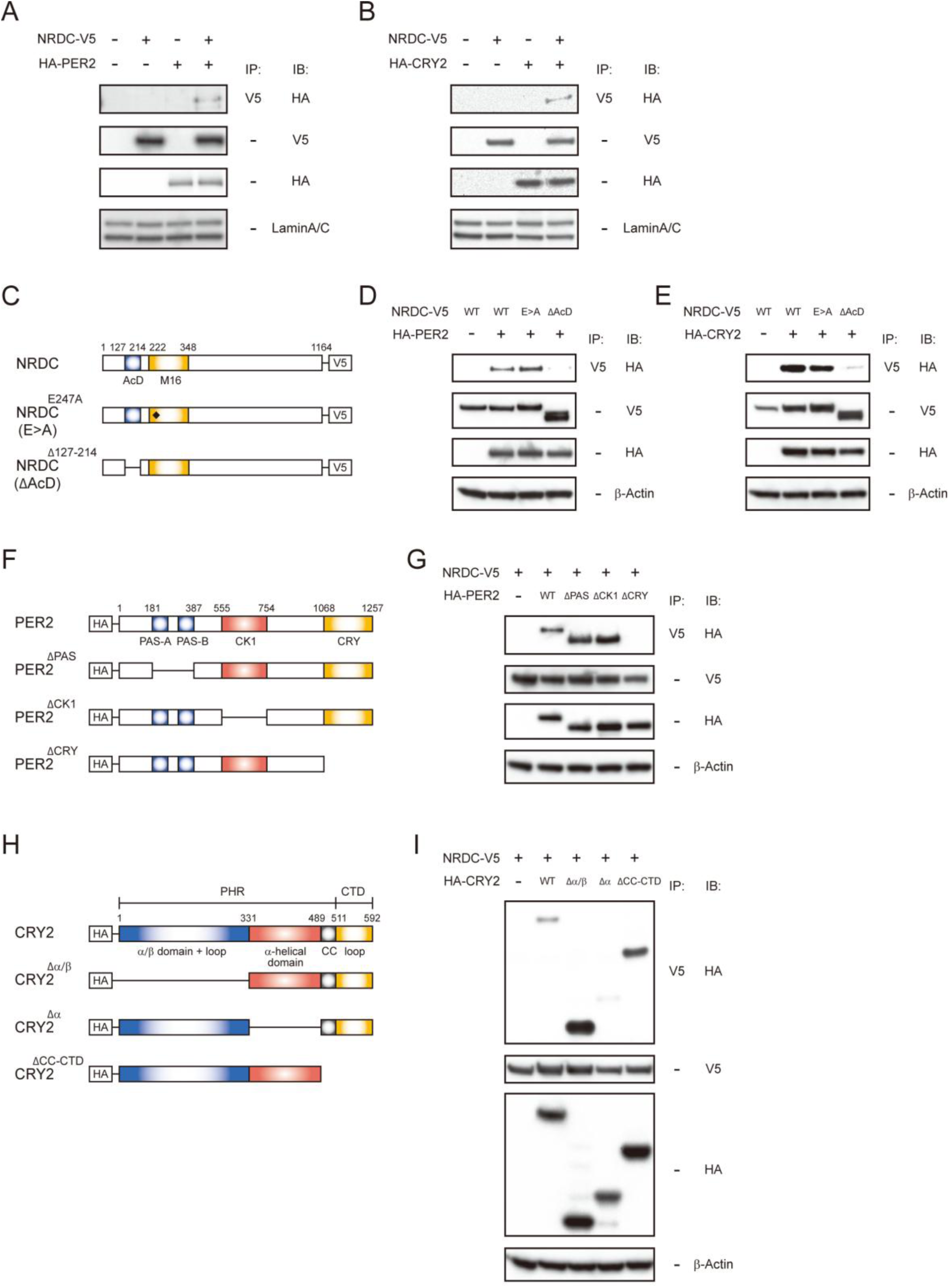
NRDC interacts with PER2 and CRY2 through defined protein domains. (A, B) Co-immunoprecipitation analysis of NRDC interaction with PER2 and CRY2 in HEK293T/17 cells, transfected with NRDC-V5 and HA-tagged PER2 (A) or CRY2 (B). Immunoprecipitates obtained with anti-V5 antibody were analyzed by immunoblotting with anti-HA antibody. (C) Schematic representation of NRDC mutants. AcD, acidic domain; ΔAcD, AcD deletion mutant; E>A, enzymatically inactive mutant. (D, E) Requirement of the NRDC acidic domain for interaction with PER2 and CRY2. HEK293T/17 cells were co-transfected with HA-PER2 or HA-CRY2 and the indicated NRDC mutants, followed by co-immunoprecipitation analysis. (F) Schematic representation of PER2 deletion mutants. PAS-A and PAS-B, PAS domain; CK1, casein kinase 1 binding domain; CRY, CRY binding domain. (G) The CRY binding domain of PER2 is required for NRDC-PER2 complex formation. Co-immunoprecipitation assays were performed as in (D). (H) Schematic representation of CRY2 deletion mutants. PHR, photolyase homology region; CTD, C-terminal tail domain; CC, coiled coil domain. (I) The α-helical domain of the CRY2 PHR is required for interaction with NRDC. Co-immunoprecipitation assays were performed as in (E).

Among M16 family metalloendopeptidases, NRDC is unique in possessing a highly acidic domain (AcD) (Fig. 2C), which has been implicated in protein-protein interactions (38). To determine whether the AcD mediates NRDC binding to PER2 and CRY2, we used an NRDC mutant lacking the AcD (ΔAcD). This deletion abolished the interaction with PER2 and CRY2, indicating that the AcD is essential for NRDC binding (Fig. 2D and E). In contrast, an enzymatically inactive NRDC mutant (E>A) retained its ability to bind to PER2 and CRY2, suggesting that the enzymatic activity of NRDC is not required for this interaction (Fig. 2D and E).

To map the NRDC-binding domain within PER2, we constructed a series of PER2 deletion mutants lacking either the tandem PER-ARNT-SIM (PAS) domain (PER2^ΔPAS^), the CK1-binding domain (PER2^ΔCK1^), or the CRY-binding domain (PER2^ΔCRY^) (Fig. 2F). Co-immunoprecipitation assays with NRDC revealed that the CRY-binding domain of PER2 is necessary for the interaction (Fig. 2G). Similarly, we mapped the NRDC-binding domain within CRY2 using three deletion mutants (Fig. 2H). Only the CRY2^Δα^ mutant, which lacks the α-helical domain, failed to bind NRDC (Fig. 2I), indicating that this region is critical for the interaction. Collectively, these results demonstrate that the AcD of NRDC, the CRY-binding domain of PER2, and the α-helical domain of CRY2 are responsible for their interaction.

### NRDC selectively modulates PER2-mediated transcriptional repression

To investigate the functional consequence of NRDC–PER2/CRY2 interactions, we performed luciferase reporter assays using an E-box-driven luciferase construct. First, we confirmed the repressive effect of PER2 and CRY2 on the CLOCK-BMAL1-mediated transcription, respectively (Fig. 3A and B). Both PER2 and CRY2 inhibited transcription in a dose-dependent manner, with CRY2 showing a stronger inhibitory effect than PER2 (Fig. 3A and B), consistent with previous reports (39).

**Fig. 3.**
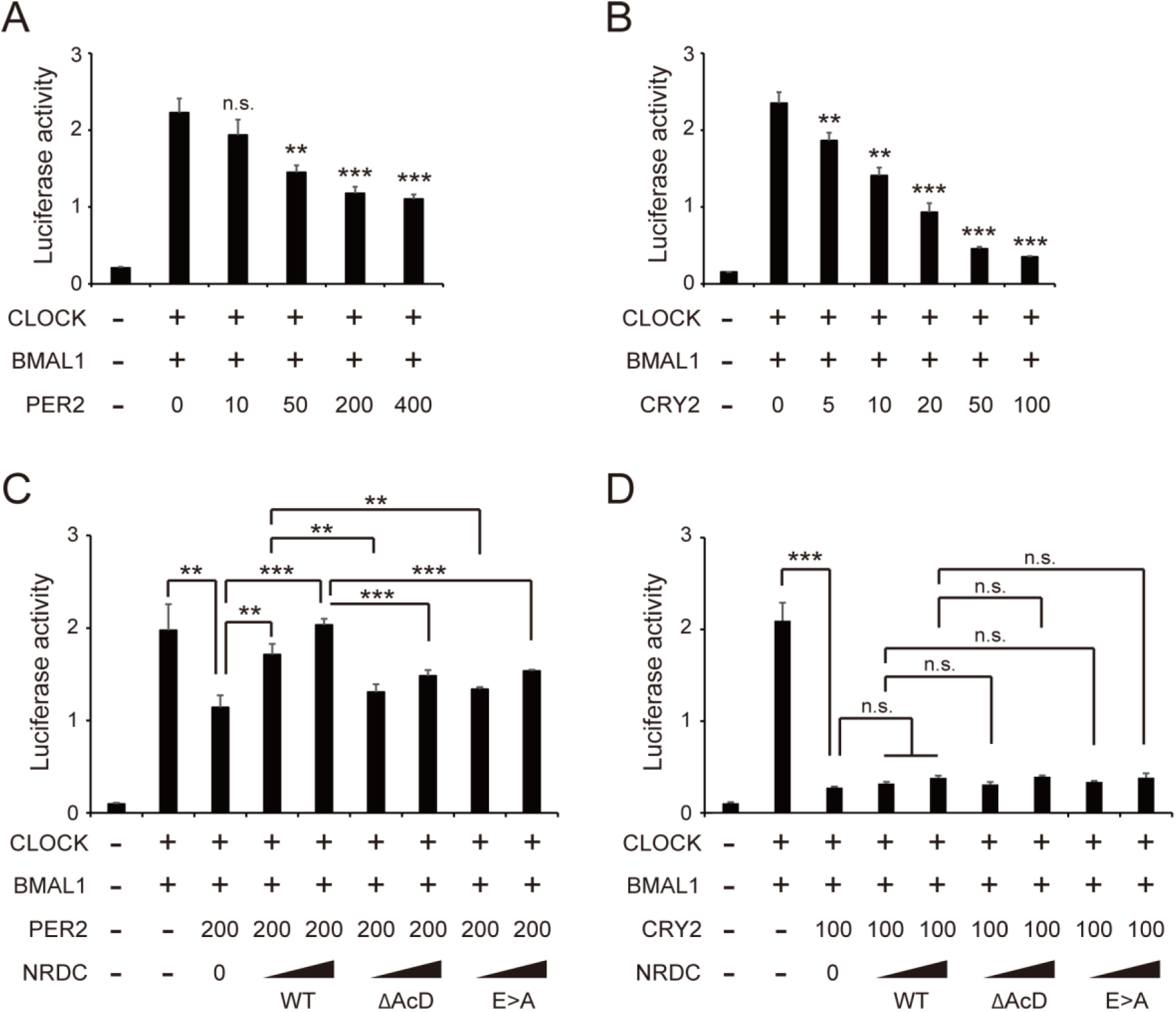
NRDC selectively modulates PER2-mediated transcriptional repression. (A, B) Dose-dependent repression of CLOCK/BMAL1-mediated transcription by PER2 and CRY2. HEK293T/17 cells were transfected with myc-CLOCK and BMAL1 expression plasmids (200 ng each) together with increasing amounts of HA-PER2 or HA-CRY2. (C, D) NRDC selectively attenuates PER2-mediated, but not CRY2-mediated transcriptional repression. HEK293T/17 cells were co-transfected with myc-CLOCK, BMAL1, and HA-PER2 or HA-CRY2 and increasing amounts of wild-type NRDC-V5 or NRDC mutants (ΔAcD or E>A). Data in (A-D) are presented as means ± SD (*n* = 3). \*\**P* < 0.01; \*\*\**P* < 0.001; n.s., not significant.

We then examined whether NRDC modulates this repression. Co-expression of NRDC significantly attenuated the inhibitory effect of PER2 (Fig. 3C), but had no effect on CRY2-mediated repression (Fig. 3D). This antagonistic effect was abolished when using either the ΔAcD or the E>A NRDC mutants, indicating that both the physical interaction through the AcD and the enzymatic activity of NRDC are required for relieving the PER2-mediated repression.

### Genome-wide identification of NRDC-binding sites in mouse liver

Our previous study showed that NRDC undergoes proteolytic processing and translocates to the nucleus, where it associates with chromatin and functions as a transcriptional coregulator in MEFs (35). In this study, we performed ChIP-seq using an anti-NRDC antibody to comprehensively identify NRDC binding sites in the mouse liver genome. From livers collected at ZT10, we identified 7,354 NRDC-binding peaks (Dataset S1). Genomic distribution analysis revealed that the majority (79.8%) of NRDC-binding sites were located in promoter regions, defined as locations within 5 kb of transcription start site (TSS) (Fig. 4A). Furthermore, NRDC signals were sharply enriched at the center of TSS (Fig. 4B), suggesting that NRDC preferentially associates with core promoter region. These findings are consistent with our previous ChIP-seq data in immortalized MEFs (35).

**Fig 4.**
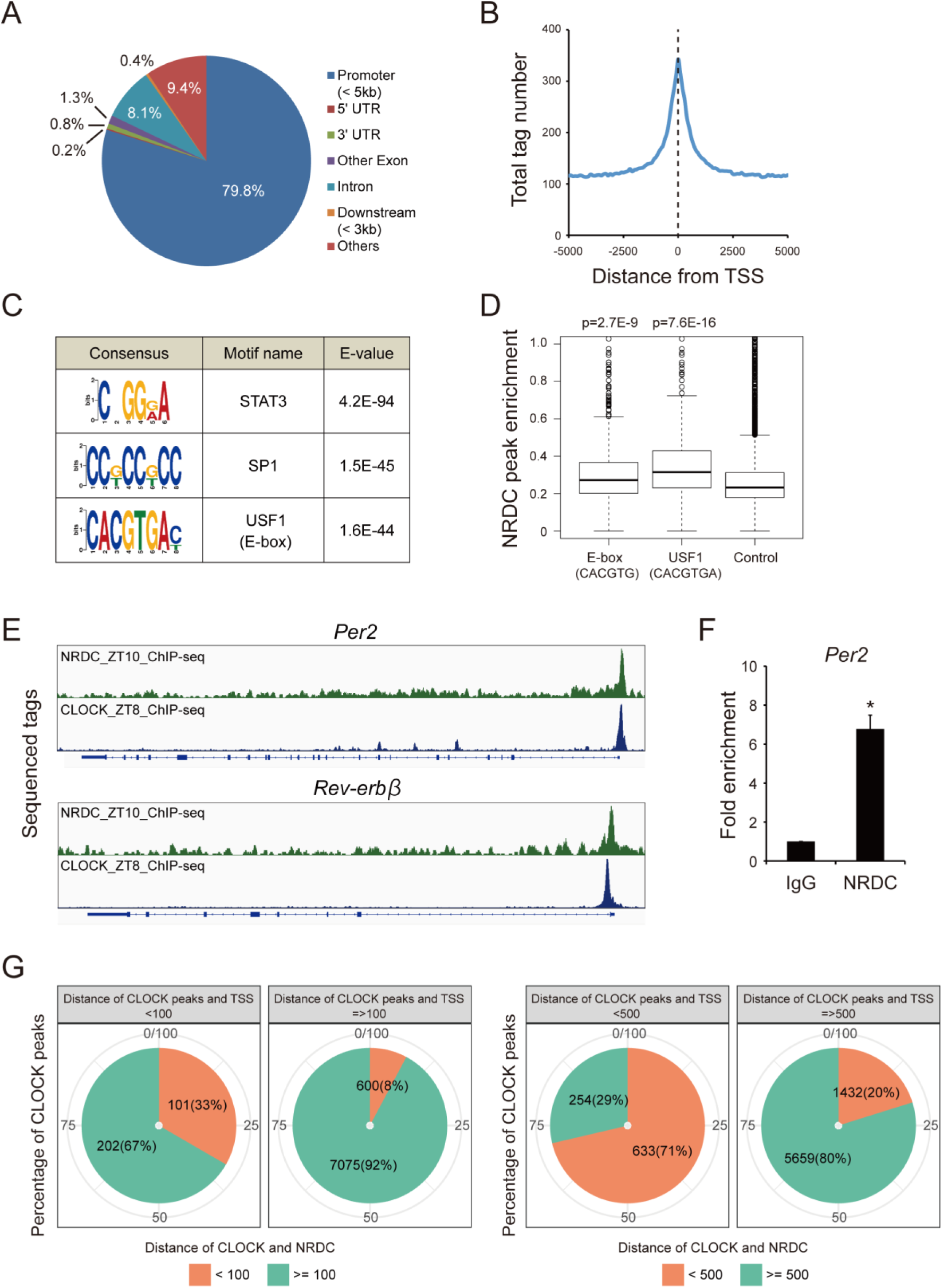
Genome wide identification of NRDC-binding sites in mouse liver by ChIP-seq. (A) Genomic distribution of NRDC ChIP-seq peaks. (B) Distribution of NRDC-binding sites relative to transcription start sites (TSSs). (C) Sequence logos of significantly enriched DNA motifs identified by MEME-ChIP. Motif names and enrichment E-values are shown. (D) ChIP-seq enrichment of NRDC at regions containing USF1 site (CACGTGA), E-box site (CACGTG), or control regions lacking both motifs. (E) Genome browser views of NRDC and CLOCK ChIP-seq signals at the *Per2* and *Rev-erbβ* loci. CLOCK ChIP-seq data were obtained from a previously published dataset (41). (F) ChIP-qPCR validation of NRDC enrichment at the E-box region of the *Per2* promoter. Chromatin was prepared from WT mice liver collected at ZT14. Enrichment was normalized to control IgG. (G) Proportions of CLOCK peaks (41) located near both TSSs and NRDC peaks (within <100 bp or <500 bp) or distant from both features.

To characterize the DNA elements associated with NRDC binding, we conducted a *de novo* motif analysis using MEME-ChIP (40). This analysis identified several enriched motifs corresponding to known transcription factors binding site, including STAT3, SP1, and USF1 (Fig. 4C). Notably, USF1 motif (CACGTGA) contains the canonical E-box motif (CACGTG), which is a well-characterized binding site for circadian transcription factors such as CLOCK and BMAL1 (1, 3). To evaluate these 2 motifs separately, we compared NRDC ChIP peak enrichment at E-box motif (CACGTG), USF1 motif (CACGTGA), and non-E-box control regions. We found that NRDC-binding peaks were significantly enriched at both E-box motifs (p=2.7E^-9^) and USF1 motifs (p=7.6E^-16^), compared with control sites (Fig. 4D). Consistently, NRDC showed marked accumulation at E-box–containing promoters such as those of *Per2* and *Nr1d2* (*Rev-erbβ*) with NRDC peaks aligning closely with CLOCK binding peaks (Fig. 4E). To validate this interaction, we performed site-specific ChIP-qPCR using primers targeting the E-box element in the *Per2* promoter, which confirmed an enrichment of NRDC at this site (Fig. 4F). Since NRDC has no known DNA-binding domain, these results suggest that NRDC is recruited to E-box motifs through interaction with circadian transcription factors. To further explore this possibility, we examined the genomic proximity of NRDC and CLOCK using previously published CLOCK ChIP-seq datasets (41). We quantified how frequently CLOCK peaks reside near both TSSs and NRDC peaks. A substantial fraction of CLOCK peaks fell in close proximity to NRDC: 33% of all CLOCK peaks were located within 100 bp of both TSSs and NRDC peaks, and this proportion increased to 71% when the window was expanded to 500 bp (Fig. 4G). In contrast, CLOCK peaks located distal to TSSs rarely overlapped with NRDC peaks (only 8% within 100 bp and 20% within 500 bp). These spatial patterns indicate that NRDC preferentially associates with CLOCK at promoter-proximal sites rather than at distal CLOCK-binding regions.

Together, our findings demonstrate that NRDC regulates PER-dependent transcriptional repression on E-box-bound CLOCK-BMAL1 complexes, thereby tuning the period length of the mammalian circadian behavioral rhythm (Fig. 5).

**Fig 5.**
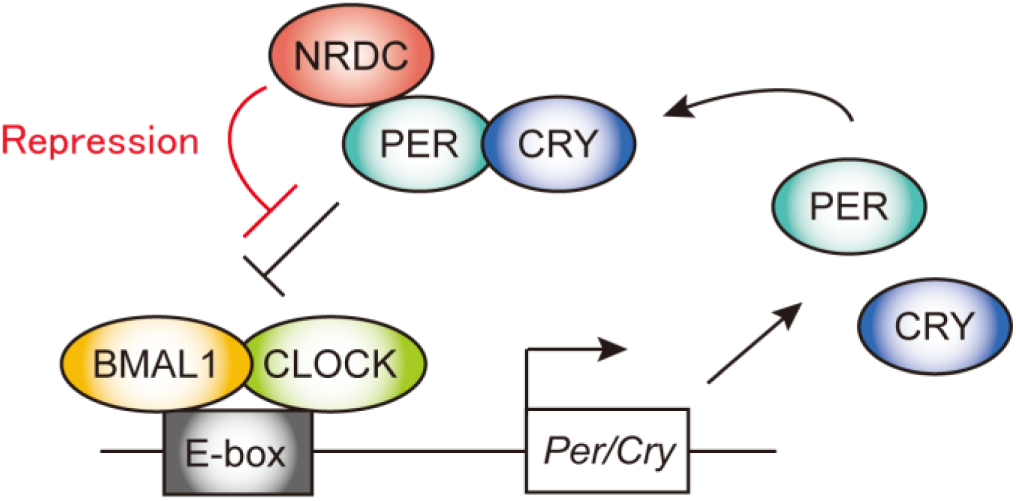
Model of the role of NRDC in the mammalian circadian clock. NRDC interacts with PER2 and regulates PER2-dependent transcriptional repression on E-box-bound CLOCK-BMAL1 complexes, thereby tuning the period length of the mammalian circadian behavioral rhythm.

## Discussion

In the present study, we identified NRDC as a novel modulator of the core circadian clock machinery through its selective interaction with PER2 and CRY2 and by antagonizing PER2-mediated transcriptional repression. NRDC-deficient mice exhibited a shorter circadian period with a significant increase in PER2 protein levels in the liver. In NRDC-KO/PER2::LUC MEFs, PER2 levels were also elevated following serum shock. Given that PER2-overexpressing mice display a similar behavioral phenotype of shortened circadian period (42), our findings suggest that NRDC regulates circadian period length, at least in part, through modulation of PER2 levels. Shortened circadian periods are relatively uncommon among genetically modified mice and have been most frequently associated with direct modification of PER or CRY stability or function (43–45). The presence of a similar phenotype in NRDC-deficient mice therefore supports the idea that NRDC acts on a regulatory step that converges on the PER2-mediated negative feedback loop, consistent with the biochemical interactions identified in this study.

We demonstrated that the interaction between NRDC and PER2/CRY2 requires the AcD of NRDC, a unique structural feature among M16 family metalloendopeptidases. Importantly, this domain was essential for the physical interaction with PER2 and CRY2. In contrast, the enzymatic activity of NRDC was dispensable for the interaction but required for the functional modulation of PER2 transcriptional activity. Our previous studies have similarly shown that the enzymatic activity of NRDC is necessary for transcriptional regulation mediated by factors such as PGC-1α (regulating UCP1) and Islet1 (regulating MafA) (32, 33). These results collectively suggest that NRDC may serve a dual role: functioning as a scaffold protein that binds circadian regulators via its AcD, and as a protease that fine-tunes their activity through enzymatic modification. Notably, NRDC selectively antagonized the transcriptional repression mediated by PER2 but not by CRY2, implying distinct regulatory pathways and binding consequences for each interaction.

Our ChIP-seq analysis of mouse liver tissue revealed significant enrichment of E-box motifs in NRDC-binding regions. Since NRDC lacks a canonical DNA-binding domain, we hypothesized that it is recruited to these sites via protein-protein interactions with transcriptional regulators. Indeed, comparison with previously published CLOCK ChIP-seq data (41) showed that over 70% of NRDC-binding peaks overlapped with CLOCK-binding regions within ±500 bp from TSS. These results support a model in which NRDC acts as a transcriptional co-regulator at E-box-containing promoters, modulating clock gene transcription in concert with core circadian components.

An important consideration raised by our findings is the possibility that subcellularly distinct pools of NRDC may be differentially regulated across the circadian cycle. NRDC is known to localize to multiple cellular compartments, including the extracellular space, cytoplasm, mitochondria and nucleus, where it exerts diverse biological functions (46). Consequently, the modest circadian variation observed in bulk NRDC levels in the liver may reflect the summation of heterogeneous NRDC populations, some of which could undergo rhythmic regulation while others remain relatively constant. Such compartment-specific oscillations could be masked in analyses of total or nuclear protein abundance. Although our data does not directly address this possibility, the multi-compartmental nature of NRDC raises the intriguing hypothesis that circadian regulation of NRDC may operate at the level of subcellular localization, interaction partners, or local proteolytic activity rather than global protein levels.

Circadian rhythms are increasingly recognized as critical regulators of systemic metabolism, immune responses, and disease susceptibility (13, 14, 47). NRDC-deficient mice have previously been reported to exhibit multiple phenotypes including hypothermia, impaired insulin secretion, and resistance to tumorigenesis and inflammation (32–34). Our findings therefore suggest that NRDC may serve as a molecular interface linking circadian timing to broader nuclear regulatory programs. Although speculative, reciprocal regulation between the circadian clock and NRDC-dependent pathways represents an attractive direction for future investigation. Elucidating how protease-based nuclear regulation intersects with circadian timing may provide new insight into the temporal organization of cellular physiology.

## Materials and Methods

### Animals

All animal experiments were performed according to procedures approved by the animal ethics committee of Kobe Gakuin University, the University of Tokyo, Shiga University of Medical Science, and Kyoto University. Mice used in this study were individually housed in cages with standard rodent chow and tap water available ad libitum under a controlled environment (temperature, 23 ± 1 °C). NRDC-deficient (NRDC-KO) mice (Accession No. CDB0466K, http://www.clst.riken.jp/arg/mutant%20mice%20list.html) were generated as previously described (29) and backcrossed to the ICR background (>98%). The NRDC-KO mice were bred with ICR-background PER2::LUC knock-in mice (36), and NRDC-KO/PER2::LUC MEF lines were prepared from these mice as previously described (12). The study is reported in accordance with ARRIVE guidelines (https://arriveguidelines.org).

### Behavioral Rhythms

Twelve- to 16-week-old male mice were housed individually in cages equipped with an area sensor and were entrained to the 12-h/12-h LD cycle for at least 2 weeks, after which they were released into DD conditions. The circadian period of the activity rhythms under the DD condition was analyzed by a χ^2^ periodogram with P < 0.001, based on the physical activity in days 1–42 after the start of DD conditions.

### Cell Culture and Plasmids for Transfection

MEFs from PER2::LUC mice (36) and NRDC-KO/PER2::LUC mice, and HEK293T/17 cells (American Type Culture Collection) were maintained at 37 °C under 5% CO2 and 95% air in Dulbecco’s Modified Eagle Medium (DMEM) (Sigma-Aldrich or Nacalai Tesuque) supplemented with 100 U/mL penicillin, 100 μg/mL streptomycin, and 10% fetal bovine serum (FBS).

The mammalian expression vectors used were Myc-CLOCK/pSG5 (a kind gift of Dr. Paolo Sassone-Corsi), BMAL1/pcDNA3.1 (a kind gift of Dr. Steven M. Reppert) and 2×HA-pcDNA3 (a kind gift of Dr. Tomohito Higashi). Full-length mouse Per2 and Cry2 cDNAs were cloned from the total cDNA of the mouse liver (C57BL/6J) by reverse transcription–PCR analysis with gene-specific primers. These cDNAs were then inserted into the 2×HA-pcDNA3 vector to generate 2×HA-PER2/pcDNA3 and 2×HA-CRY2/pcDNA3. Deletion mutants of PER2 (PER2^ΔPAS^: the PAS domain deletion mutant of PER2, PER2^ΔCK1^: the CK1 binding domain deletion mutant of PER2, PER2^ΔCRY^: the CRY binding domain deletion mutant of PER2) and CRY2 (CRY2^Δα/β^: the mutant CRY2 lacking the α/β domain and the inter-loop domain of the PHR, CRY2^Δα^: the mutant CRY2 lacking the α-helical domain of the PHR, CRY2^ΔCC-CTD^: the mutant CRY2 lacking the CC and the CTD) were generated by using the PCR technique. The primers are listed in Supplementary Table 1. mNRDC-V5/pcDNA3.1 and the mutants of NRDC (NRDC^ΔAcD^ and NRDC^E>A^) were obtained as described previously (33). Transfections were carried out using X-tremeGENE HP (Roche) according to the manufacturer’s instructions.

### Real-Time Monitoring of Cellular Rhythms

Real-time monitoring of the cellular bioluminescence rhythms was performed as described previously (48) with minor modifications. In brief, PER2::LUC MEFs were plated on 35-mm dishes (1.0 × 10^6^ cells/dish) and then cultured at 37 °C under 5% CO2. After 24 h, the cells were treated with 0.1 μM (final concentration) dexamethasone for 2 h, after which the media were replaced by recording media [phenol-red free DMEM (Sigma-Aldrich) supplemented with 10% FBS, 3.5 g/L glucose, 25 U/mL penicillin, 25 μg/mL streptomycin, 0.1 mM luciferin, and 10 mM Hepes-NaOH, pH 7.0]. The bioluminescence signals of the cultured cells were continuously recorded for 5–10 days at 37 °C in air with a Kronos AB-2500 or AB-2550 (Atto) luminometer.

### Preparation of Nuclear Proteins from Mouse Tissue

The nuclear proteins and cytoplasmic proteins were isolated as described previously (49). In brief, the mouse tissue (1 g wet weight) was washed with ice-cold PBS and homogenized at 4 °C with 9 mL of ice-cold buffer A composed of 10 mM Hepes-NaOH (pH 7.8), 10 mM KCl, 0.1 mM EDTA, 1 mM DTT, 1 mM phenylmethylsulfonyl fluoride (PMSF), 4 μg/mL aprotinin, 4 μg/mL leupeptin, 50 mM NaF, and 1 mM Na3VO4. The homogenate was centrifuged twice (5 min each, 700 × g), and the resultant precipitate was resuspended in 2 mL of ice-cold buffer C composed of 20 mM Hepes-NaOH (pH 7.8), 400 mM NaCl, 1 mM EDTA, 5 mM MgCl2, 2% (vol/vol) glycerol, 1 mM DTT, 1 mM PMSF, 4 μg/mL aprotinin, 4 μg/mL leupeptin, 50 mM NaF, and 1 mM Na3VO4. After gentle mixing at 4 °C for 30 min, the suspension was centrifuged twice (30 min each, 21,600 × g), and the final supernatant was used as “nuclear extract.”.

### Antibodies and Immunoblot Analysis

Rat anti-mouse NRDC monoclonal antibody (clone #135) and mouse anti-mouse NRDC monoclonal antibody (clone #2E6), commercially available at Merck Millipore (MABS2057 & 2058), were originally raised in our laboratory (32, 35). Other antibodies were from the following sources; anti-CLOCK (clone CLSP3: MBL) (49), anti-BMAL1 (clone BIBH2: MBL) (49), anti-PER2 (ADI), anti-CRY1 (MBL), anti-CRY2 (MBL), anti-DBP (MBL), anti-E4BP4 (MBL), anti-HA (Roche), anti-V5 (Invitrogen), anti-Lamin A/C (Santa Cruz) and anti-β-actin (Santa Cruz).

Preparations of total cell extract and immunoblot experiment were carried out as described previously (32). In brief, cells were lysed in the buffer containing 10 mM Tris–HCl pH 7.4, 150 mM NaCl, 1% NP-40, protease inhibitor cocktail (Roche) and phosphatase inhibitor cocktail (Sigma). Cell lysates were separated by SDS–polyacrylamide gel electrophoresis and transferred onto polyvinylidene difluoride (PVDF) membranes (Millipore). After blocking, membranes were incubated with primary antibodies, followed by horseradish peroxidase-conjugated secondary antibodies. The immobilized peroxidase activity was detected with the enhanced chemiluminescence system (Millipore).

For isolation of nuclei, cells were suspended in hypotonic buffer containing 10 mM HEPES (pH 7.9), 1.5 mM MgCl2, 10 mM KCl, 0.1 mM EDTA, 0.1% NP-40, 1 mM dithiothreitol, protease inhibitor cocktail (Roche) and phosphatase inhibitor cocktail (Sigma), followed by homogenization and centrifugation (3,000 rpm, 5 min). Isolated nuclei were then resuspended in high salt buffer containing 20 mM HEPES (pH 7.9), 1.5 mM MgCl2, 400 mM NaCl, 0.1 mM EDTA, 10% glycerol, 0.1% NP-40, 1 mM dithiothreitol, protease inhibitor cocktail (Roche) and phosphatase inhibitor cocktail (Sigma), followed by centrifugation (14,000 rpm, 5 min).

### Dual-Glo Luciferase Reporter Assay

HEK293T/17 cells in 12-well plates were transiently transfected with 200 ng of Myc-CLOCK/pSG5, 200 ng of BMAL1/pcDNA3.1, 20 ng of a firefly luciferase plasmid AVP-E-box SV40-Luc/pGL3, and a combination of various expression plasmids using X-tremeGENE HP. A promoter null mutant (Δ27-779 nucleotides) renilla luciferase plasmid pGL4.74[*hRluc*/TK]-promoter null (100 ng/well, Promega) was used as an internal control. The total amount of DNA was adjusted by adding the empty expression plasmids. The transfected cells were harvested 36 h after the transfection, and the cell extracts were subjected to Dual-Glo luciferase assays according to the manufacturer’s protocol (Promega).

A single set of the following oligonucleotides was designed to generate the promoter null mutant: sense; 5’-CCGGTACCTGAGTCTAAGCTTGGCAATCCG-3’, and antisense; 5’-CGGATTGCCAAGCTTAGACTCAGGTACCGG-3’. The promoter null mutant was generated by using the PCR technique, and the mutations were confirmed by DNA sequencing.

### Chromatin Immunoprecipitation (ChIP), ChIP-Sequencing and data processing

ChIP with anti-mouse NRDC antibody (clone #2E6) or control IgG was performed using the ChIP-IT Express kit (Active motif) according to the manufacturer’s protocol (35). ChIP-seq library preparation and sequencing were outsourced to Active Motif (Carlsbad, CA). Briefly, sequencing was carried out on an Illumina NextSeq 500 platform generating 75-bp single-end reads. After removal of reads that did not pass the Illumina purity filter, sequences were aligned to the reference genome using BWA (default parameters). Only uniquely mapped reads with ≤2 mismatches were retained, and PCR duplicates were removed. Fragment density profiles were generated by extending the 5’ ends of aligned reads in silico to the average fragment length (150–250 bp), followed by binning into 32-nt windows to compute genome-wide signal maps. Peak calling was performed using MACS for transcription factor–like sharp peaks and SICER for broad enrichment regions, using matched Input DNA as background (Dataset S1, “NRDC-liver-Intervals”). Annotation of Intervals and Active Regions— including genomic features, CpG islands, promoters, and nearest genes—was performed using Active Motif’s analysis pipeline. The raw FASTQ files have been deposited in GEO (GSE314983).

### Statistical Analysis

Data are presented as mean values ± the standard deviation (SD) or standard error (SE). Differences between groups and genotypes were analysed using an unpaired 2-tailed Student’s t-test for 2 groups or a one-way ANOVA (the post hoc Tukey–Kramer HSD test) for more than 3 groups. The relationship of the relative enrichment of read densities among promoter regions between each ChIP-seq experiment was assessed using Pearson’s correlation coefficient r. The Wilcoxon rank-sum test was used to calculate the significance of differences in enrichment.

## Acknowledgments

This study was supported by Grants-in-Aid for Scientific Research (KAKENHI: 16K08536, 19K05958, 23K07550, 23K07966 and 25K02686) from Japan Society for the Promotion of Science (JSPS). It was also supported by the Takeda Science Foundation, the Kurata Grants by the Hitachi Global Foundation, Kao Research Council for the Study of Healthcare Science, the NOVARTIS Foundation (Japan) for the Promotion of Science and Manpei Suzuki Diabetes Foundation. We are grateful to T. Kita for continuous encouragement.

## Data Availability

ChIP-seq data obtained in this study have been deposited in NCBI Gene Expression Omnibus (GEO) under accession number GSE314983. Any additional information required to reanalyse the data reported in this paper is available upon reasonable request.

## Author Contributions

Y.H., Y.F., H.Y., and E.N. designed the research; Y.H., R.N., M.O., K.N., and H.Y. performed the research; Y.H., R.N., M.O., Y.M., Y.K., K.N., and H.Y. analyzed the data; Y.F. and E.N. supervised the project; Y.H. and H.Y. wrote the manuscript; and N.K., Y.F. and E.N. reviewed and edited the manuscript.

